# Deletion of miR-146a enhances therapeutic protein restoration in model of dystrophin exon skipping

**DOI:** 10.1101/2023.05.09.540042

**Authors:** Nikki M. McCormack, Kelsey A. Calabrese, Christina M. Sun, Christopher B. Tully, Christopher R. Heier, Alyson A. Fiorillo

**Author notes:** **Correspondence should be addressed to:** Alyson A. Fiorillo, PhD Center for Genetic Medicine Research, Children’s Research Institute 7144 13th Place NW, Washington DC, 20012.

## Abstract

Duchenne muscular dystrophy (DMD) is a progressive muscle disease caused by the absence of dystrophin protein. One current DMD therapeutic strategy, exon skipping, produces a truncated dystrophin isoform using phosphorodiamidate morpholino oligomers (PMOs). However, the potential of exon skipping therapeutics has not been fully realized as increases in dystrophin protein have been minimal in clinical trials. Here, we investigate how miR-146a-5p, which is highly elevated in dystrophic muscle, impacts dystrophin protein levels. We find inflammation strongly induces miR-146a in dystrophic, but not wild-type myotubes. Bioinformatics analysis reveals that the dystrophin 3′UTR harbors a miR-146a binding site, and subsequent luciferase assays demonstrate miR-146a binding inhibits dystrophin translation. In dystrophin-null *mdx52* mice, co-injection of miR-146a reduces dystrophin restoration by an exon 51 skipping PMO. To directly investigate how miR-146a impacts therapeutic dystrophin rescue, we generated *mdx52* with body-wide miR-146a deletion (*146aX*). Administration of an exon skipping PMO via intramuscular or intravenous injection markedly increases dystrophin protein levels in *146aX* versus *mdx52* muscles; skipped dystrophin transcript levels are unchanged, suggesting a post-transcriptional mechanism-of-action. Together, these data show that miR-146a expression opposes therapeutic dystrophin restoration, suggesting miR-146a inhibition warrants further research as a potential DMD exon skipping co-therapy.

## Introduction

Duchenne muscular dystrophy (DMD) is a progressive muscle disease caused by out-of-frame mutations in the dystrophin gene resulting in the complete loss of dystrophin protein^1^. DMD is characterized by severe muscle weakness and chronic muscle inflammation leading to loss of ambulation and a shortened lifespan. Becker muscular dystrophy (BMD) is an allelic disease where the reading frame is preserved, resulting in production of a truncated dystrophin isoform that is expressed at lower and more variable levels^2–4^. BMD is less severe and shows high variability both in clinical presentation and in levels of dystrophin protein even when comparing subjects harboring the same in-frame mutation^5–8^. Interestingly, while dystrophin protein levels are variable and lower-than-normal in BMD, there are no differences in the amount of dystrophin mRNA detected in BMD as compared to unaffected/healthy muscles^7, 8^.

Precision medicine strategies for DMD such as gene therapies and exon skipping antisense oligonucleotides (AOs) seek to convert a severe DMD genotype (dystrophin null) into a milder BMD (truncated dystrophin) phenotype. Specifically, exon skipping AOs act by binding to a complementary sequence on the dystrophin pre-mRNA to promote the exclusion of the frame-shifting exon. Currently four exon skipping phosphorodiamidate morpholino oligomers (PMOs) have received accelerated FDA approval; these target dystrophin exons 51 (Eteplirsen), 53 (Golodirsen, Viltepso) and 45 (Casimersen)^9–12^. However, results from clinical trial data show low and variable dystrophin rescue. As an example, for the first approved exon skipping drug (Eteplirsen), the Western blot analysis showed an average of 0.93% dystrophin in treated muscles compared with a baseline value of 0.08% in biopsies of untreated Duchenne muscles^13^. In the later Golodirsen clinical trial, mean dystrophin protein levels measured via Western blot were 1.019% of normal dystrophin levels despite showing greater levels of skipped dystrophin transcripts (16.363%)^10, 14^.

The disconnect between the extent of dystrophin exon skipping (mRNA) and dystrophin protein levels point to post-transcriptional factors in the muscle that affect total dystrophin protein production. microRNAs (miRNAs) are elevated in a variety of muscle diseases; their canonical role is to fine-tune gene expression by binding to the 3′UTR of target genes and most often function to regulate protein levels post-transcriptionally via translation inhibition^7, 15–17^. Previously, we identified several miRNAs that bind the dystrophin 3’UTR and inhibit dystrophin protein production (termed dystrophin targeting miRNAs or DTMs)^7^. These include miR-146a-5p, which is elevated in DMD and BMD muscle biopsies and is induced by inflammation *in vitro*, suggesting regulation by the pro-inflammatory transcription factor NF-κB. Indeed, others have demonstrated NF-κB-specific regulation of miR-146a using cell culture studies^18^ and bioinformatic analysis from ENCODE data^19^. Importantly, our lab previously found that levels of miR-146a were inversely associated with dystrophin restoration in the muscles of DMD model (*mdx)* mice treated with an exon skipping PMO^7^. In a separate study, we identified a panel of 9 inflammatory miRNAs that are elevated in the *mdx* diaphragm and are reduced by treatment with the corticosteroid prednisone or the dissociative steroid vamorolone; included in this panel is the DTM miR-146a^20^. Finally, we recently reported that treating BMD model (*bmx*) mice with prednisone or vamorolone dampens expression of DTMs including miR-146a and increases dystrophin protein levels in the gastrocnemius and heart of bmx mice by ∼50%^21^. Collectively, these data suggest that miR-146a is regulated by inflammation, is elevated in dystrophic muscle, and inhibits dystrophin protein production.

In the present study, we hypothesized that genetic deletion of miR-146a specifically would improve dystrophin restoration in dystrophin-deficient mice harboring a deletion of *Dmd* exon 52 (*mdx52*). To test this, we generated double knockout mice and carried out a proof-of-concept study, performing high dose systemic administration of an exon 51 skipping PMO to *mdx52* and *mdx52*;*miR-146a*^-/-^ (*146aX*) mice, where our primary readout was extent of dystrophin protein restoration. Excitingly, following both intramuscular and systemic delivery of PMO, we show deletion of miR-146a significantly increases dystrophin protein levels.

## Methods

### Animal care and maintenance

All animal studies were done in adherence to the NIH Guide for the Care and Use of Laboratory Animals. All experiments were conducted according to protocols within the guidelines and under approval of the Institutional Animal Care and Use Committee of Children’s National Hospital. *mdx52* mice contain a deletion of exon 52 of the *Dmd* gene, resulting in the absence of full-length dystrophin. These mice on a C57/BL6 background are maintained in our own facility and were originally a kind gift from Dr. Shin’ichi Takeda. Male C57/BL6 mice were purchased from JAX laboratory (Bar Harbor, ME). *bmx* mice contain an endogenous deletion of murine dystrophin exons 45-47, are maintained on a C57/BL6J background, and were previously characterized ^8^.

### Muscle biopsies

Human biopsies used here were banked diagnostic biopsies obtained from the vastus lateralis muscle. Six BMD muscle biopsies were obtained from the Eurobiobank, Telethon Network Genetic Biobanks (GTB 12001D) (ID #: 2518, 3010, 3419, 4228, 7560, 7827). The remaining 4 BMD biopsies were obtained from our muscle biopsy bank as previously reported (ID#: 1141, 3212, 3255, 3461)^4, 7^.

### Generation of mdx/146a^-/-^ mice

*mdx52* mice (*dmd^-/-^* or *dmd^-/Y^*) on a C57/BL/6 background are bred in house^22^. *miR-146a-/-* (B6.Cg-*Mir146^tm^*^1^*^.1Bal^*/J) have been previously described^23^ and were purchased from The Jackson Laboratory (Strain # 016239) and are on a C57/BL/6J background. To generate *mdx52* mice with deletion of miR-146a, dmd^-/-^;146a^+/+^ females were crossed with dmd^+/Y^;146a^-/-^ males to generate dmd^-/+^:146a^+/-^ females and dmd^-/Y^;146a^+/-^ males. dmd^-/+^;146a^+/-^ females and dmd^-/Y^;146a^+/-^ males were then crossed to generate dmd^-/-^;146a^-/-^ and dmd^-/Y^;146a^-/-^ breeders. Breeders were crossed to generate dmd^-/Y^;146a^-/-^ mice (*146aX*). For all experiments only males were utilized as DMD is an X-linked disorder. Genotyping for *mdx52* and miR-146a KO alleles was performed using TransnetYX with custom primers in *Dmd* exon 3 and a neomycin cassette for *mdx52* and with the standard JAX protocol for miR-146a^-/-^ with forward primer 24664 (5′-GCT TAT GAA CTT GCC TAT CTT GTG-3′) and reverse primer 24665 (5′-CAG CAG TTC CAC GCT TCA-3′).

### Cell culture and treatment of H2K myotubes

H2K (H2Kb-tsA58) WT and *mdx* myoblasts were cultured as previously described^24^. H2K *mdx* or WT myoblasts were differentiated into myotubes in 12-well plates (1.25 × 105 cells/well) with Matrigel at 37°C. For TNF-α inductions, after 4 days of differentiation, myotubes were induced with TNF-α (10 ng/ml) for 24 hr. After 24 h, cells were lysed for RNA using TRIzol and expression of miR-146a and control RNAs was quantified by TaqMan Assay (Thermo Fisher Scientific). Comparison of miR-146a induction between groups was performed using a Two-Way ANOVA with a Holms-Sidak post-hoc analysis. For LPS-treatment, seeded myoblasts were differentiated into myotubes in 12-well plates (1.25 × 105 cells/well) with Matrigel at 37°C. After 4 days of differentiation, myotubes were then treated with LPS to induce inflammation (Thermo Fisher Scientific) at a dilution of 1:1,000. After 24 h, cells were lysed for RNA using TRIzol and expression of miR-146a and control RNAs was quantified by TaqMan Assay (Thermo Fisher Scientific).

### Cloning

Cloning was performed as previously reported^7^. Briefly, the entire 2.7kb of the dystrophin 3′UTR was amplified from normal muscle using the Flex cDNA synthesis kit (Quanta) using primers that added 5′ XhoI and 3′ NotI restriction enzyme sequences (Forward primer “017 DMD 3′UTR”- 5′CCGCTCGAGGCG AAGTCTTTTCCACATGGCAGAT3′ Reverse primer “018 DMD 3′UTR”- 5′CGGCGGCCGC GTAACATAACTGCGTGCTTTATT3′. The amplified product was gel purified, digested with XhoI and NotI, and ligated into the pSiCheck2 plasmid (Promega Cat #C8021) where renilla luciferase is upstream of the 3’UTR and firefly luciferase is driven by a separate promoter as a reference control. Ligation products were transformed into TOP10 cells (Life Technologies) using ampicillin selection. Positive clones were verified by diagnostic digests and Sanger sequencing.

#### Luciferase in vitro Reporter Assay

HEK293 cells were seeded in 24-well plates at a density of 4 × 104 or 8 × 10^4^ cells/well and co-transfected 24 hours later with 200 ng of the PsiCheck2 plasmid (Promega, cat #C8021) with the human dystrophin 3′ UTR or mutant 3′ UTR and with 50 nM miR-146a mimic (Life Technologies) with Lipofectamine 2000 according to the manufacturer’s protocol. Cells were harvested 24 h later according to Dual-Glow Luciferase Reporter Assay System protocol (Promega) where renilla luciferase was the main readout (3′UTR) and firefly luciferase served as the internal control.

### miR-146a mimic in vivo injections

*mdx52* mice were anesthetized with isoflurane and 2 μg of PMO was injected into each tibialis anterior. Mice were co-injected with 10 μg of either a miR-146a mimic or a control miRNA sequence (ThermoFisher scientific). Mice were euthanized two weeks after intramuscular injection and muscles were harvested.

### PMO treatment of dystrophic mice

The *Dmd* exon 51 skipping PMO utilized here is the sequence for Eteplirsen modified for the mouse genome and has been previously reported ^25^. The sequence is: (5′-CTCCAACAGCAAAGAAGATGGCATTTCTAG-3′) and was synthesized by Gene Tools. For intramuscular injections, 12- week-old mice were anesthetized with isoflurane and 2 μg of PMO was administered into each tibialis anterior. Mice were euthanized two weeks after intramuscular injection and muscles were harvested and frozen.

For intravenous injections, 15-week-old mice were anesthetized with isoflurane and 400 mg/kg PMO was administered via retroorbital injection. A second IV dose of PMO (400 mg/kg) was given seven days later. Mice were euthanized two weeks after the second injection and muscles were harvested and frozen.

### Capillary electrophoresis (Wes)

Muscles were dissected and frozen in liquid-nitrogen cooled isopentane. 8 µm sections were lysed in 5% SDS buffer containing 10mM EDTA (pH 8.0), 75mM Tris-HCL (pH 6.8), and protease inhibitors. Capillary Western immunoassay (Wes) analysis was performed according to manufacturer’s instructions using 66–440 kDa Separation Modules (ProteinSimple). In each capillary 0.2mg/mL protein was loaded for analysis with antibodies to dystrophin (Abcam # ab15277, dilution 1:15) as in ^26^, or vinculin (Abcam #ab130007, dilution 1:100), and anti-rabbit secondary (ProteinSimple #042-206). Compass for SW software was used to quantify chemiluminescence data.

### qRT-PCR

#### Dmd exon skipping

Total RNA was extracted from muscle samples by standard TRIzol (Life Technologies) isolation. RNA quantity and quality was determined using a nanodrop 2000. 400□ng of RNA was reverse-transcribed using Random Hexamers and High-Capacity cDNA Reverse Transcription Kit (Thermo Fisher). Skipped and non-skipped *Dmd* transcripts were measured by qRT-PCR using TaqMan assays and Taqman Fast Advanced MasterMix (Thermo Fisher) on an ABI QuantStudio 7 Real-Time PCR machine (Applied Biosystems). A custom TaqMan probe for the skipped *Dmd* product (Thermo Fisher) was designed to amplify the splice junction spanning *Dmd* exons 50-53. TaqMan probe Mm01216492_m1 (Thermo Fisher) that amplifies the region spanning *Dmd* exons 2–3 was utilized to calculate non-skipped *Dmd* transcript levels. Percent exon skipping was calculated based on the corresponding ΔCt values for skipped and total *Dmd* transcript normalized to *Hprt* (Mm01545399_m1; Thermo Fisher) and *18S* (Mm03928990_g1; Thermo Fisher) mRNA, using the following equation: [(Average of triplicate reactions of skipped *Dmd*)/(Average of triplicate reactions of skipped *Dmd*□+□Average of triplicate reactions of non-skipped *Dmd*)] * 100.

#### miRNAs

Total RNA was converted to cDNA using multiplexed RT primers and High Capacity cDNA Reverse Transcription Kit (ThermoFisher; Carlsbad, CA). miRNAs were then quantified using individual TaqMan assays on an ABI QuantStudio 7 real time PCR machine (Applied Biosystems). Assay IDs include: miR-146a 000468 and controls sno202 001232, RNU48 001006. miR-146a levels were normalized to the geometric mean of the two control genes^27^.

*All assay IDs are listed below in Table 1:*

| Oligonucleotide Name | Sequence | Purpose |
| --- | --- | --- |
| Dmd exons 50-53 | Thermo Fisher Scientific Custom Assay ID ARKA43N | qRT-PCR |
| Dmd exons 2-3 | Thermo Fisher Scientific Assay ID Mm01216492_m1 | qRT-PCR |
| Hprt | Thermo Fisher Scientific Assay ID Mm01545399_m1 | qRT-PCR |
| 18S | Thermo Fisher Scientific Assay ID Mm03928990_g1 | qRT-PCR |
| miR-146a | Thermo Fisher Scientific Assay ID 000468 | qRT-PCR |
| sno202 | Thermo Fisher Scientific Assay ID 001232 | qRT-PCR |
| RNU48 | Thermo Fisher Scientific Assay ID 001006 | qRT-PCR |

### Immunofluorescence of muscle samples

Muscles were mounted on cork, flash frozen, and sectioned (8 µm) onto slides. For dystrophin and F4/80 immunofluorescence experiments muscle sections were fixed in ice cold acetone for 10 minutes. Slides were washed with 1X PBST (0.1% Tween 20), blocked for 1 hour (1X PBST 230 with 0.1% Triton X-100, 1% BSA, 10% goat serum, and 10% horse serum), washed 3 times, then exposed to primary antibodies overnight at 4°C. For dystrophin immunostaining these included: anti-dystrophin 1:150 (Abcam, #ab154168) as in^28^ and rat anti-laminin-2 1:100 233 (Sigma-Aldrich, Cat. #L0663). For macrophage staining these included: anti-F4/80 which recognizes the mouse F4/80 antigen, a 160kD glycoprotein expressed by murine macrophages 235 (1:100, Bio-Rad clone C1:A3-1, Cat. MCA497) and rabbit anti-laminin (1:50, Sigma-Aldrich Cat. 236 #L9393). Secondary antibodies included: goat anti-rabbit 568 1:400 (ThermoFisher, #A-11036), 237 donkey anti-rat 488 1:400 (ThermoFisher, #A-21208), and goat anti-rabbit 647 1:400 238 (ThermoFisher, #A-21245). Wheat germ agglutinin (WGA) conjugated to Alexa Fluor 647 (Life Technologies, MA) was prepared as 1 mg/ml stock solutions and used at a 1:500 dilution in PBS. Coverslips were mounted using Prolong Gold Mounting Medium 239 with DAPI. Slides were imaged using an Olympus VS-120 scanning microscope at 20X.

### Dystrophin immunofluorescence image analysis

Images in each experiment were thresholded to the same levels, and then exported as PNG files and blinded using randomly assigned number IDs. To determine % dystrophin-positive fibers, the total number of myofibers (marked by either wheat-germ agglutinin or laminin staining) were counted manually using ImageJ. Dystrophin positive fibers were additionally counted, and the percentage of dystrophin-positive fibers were determined by the following formula: dystrophin-positive fibers/ total fibers x 100. For secondary analysis dystrophin high and low fiber percentages were determined based on a visual standard (refer to **Figure 6C**).

### Histological staining

Hematoxylin and eosin was performed to analyze necrosis and inflammation. Briefly, tissue sections were stained for 10 minutes with hematoxylin and excess stain was removed in running tap water until nuclei turned blue. Slides were incubated in 70% ethanol for 3 minutes and then stained with eosin for 3 minutes. Tissue sections were dehydrated with ethanol and cleared with xylene. Coverslips were mounted with permount. All slides were imaged using an Olympus VS-120 scanning microscope at 20X.

### Muscle histology analysis

Cross-sectional area (CSA) and minimum Feret’s diameter were determined using the MuscleJ macro for Fiji.^62^ The variance coefficients for CSA and minimum Feret’s diameter were calculated by dividing standard deviation by the average and multiplying by 1000. Hematoxylin and Eosin staining was performed to assay necrosis and inflammation. ImageJ was used to outline areas of necrosis and inflammation.

### Motor function tests

Motor function tests were performed in PMO treated mice 1 week after second injection (17 weeks of age). Forelimb and hindlimb grip strength was assessed using a grip strength meter (Columbus Instruments) daily for 2 consecutive days according to Treat NMD protocols (DMD_M.2.2.001), with data interpreted as average maximum daily values. Two-limb wire hang and four-limb grid hang tests were performed in accordance with Treat NMD protocols (DMD_M.2.1.005). For two-limb wire hang, a wire hanger was suspended placed ∼35cm above a cage with soft bedding. Mice were hung using only their forelimbs; however, they were allowed to swing and hang with all four limbs if able. Hang time was recorded, with 600 seconds used as a cutoff.

### Statistical Analysis

For assays with greater than two groups, measurements were compared between groups using one-way ANOVA unless otherwise indicated. Post hoc linear tests between each group were performed; the resulting p value reported in figures was adjusted for multiple testing via the Holmes-Sidak method. The contrasting groups in all post hoc comparisons are indicated in each figure. For assays with two groups where the null hypothesis was testing a change in one direction, a one-tailed, Student’s t test was utilized, whereas in assays where the potential change between groups was + or −, a two-tailed Student’s t test was utilized. For all bar graphs, data are presented as mean ± standard error of the mean (S.E.M). Details of statistical tests are specified in the figure legends.

## Results

### Inflammation-regulated miR-146a is elevated in dystrophic muscle

We previously reported that DTMs, including miR-146a, inhibit dystrophin protein production *in vitro* and are inversely related to exon skipping-mediated dystrophin restoration *in vivo*^7^. One particular miRNA, miR-146a, was found to be highly elevated in BMD, DMD and *mdx* muscle^7^ and analysis of ENCODE data shows it is regulated by the pro-inflammatory transcription factor NF-κB^19^.

The miR-146a locus is found on chromosome 5q33.3 and is located in a cluster along with the long non-coding RNA MIR3142HG. miR-146a is evolutionarily conserved across species, as demonstrated by alignment of the miR-146a sequence from human, chimpanzee, baboon, dog, mouse, xenopus and zebrafish (**Figure 1A**). Evaluating levels of miR-146a from Becker muscular dystrophy human biopsies with deletion of *DMD* exons 45-47, we previously found miR-146a levels are higher in biopsies from patients with low dystrophin levels (less than 20% of unaffected control biopsies, **Figure 1B**, data from^7^.) Plotting miR-146a levels versus dystrophin protein levels (as measured by Western blot^7^) in these same BMD 45-47 biopsies, we observed a significant inverse correlation between miR-146a levels and the percent of dystrophin protein in BMD biopsies (Spearman correlation, r=-0.6365, p<0.01, **Figure 1B).** Elevated miR-146a levels in dystrophic muscle are corroborated by our analysis of previously published data from Eisenberg et. al^15^ showing miR-146a is upregulated in DMD, and BMD patient biopsies as compared to unaffected control biopsies (**Figure 1C**). We analyzed the diaphragm, tibialis anterior, quadriceps, gastrocnemius, and triceps muscles from 3-month-old *mdx52* mice and found elevated levels of miR-146a in all muscles analyzed (**Figure 1D**). We also analyzed miR-146a in the *bmx* mouse model of BMD^8^ with deletion of murine *Dmd* exon 45-47 at 5 months of age, and saw significantly elevated levels in the quadriceps, gastrocnemius and triceps (**Figure 1E**).

**Figure 1.**
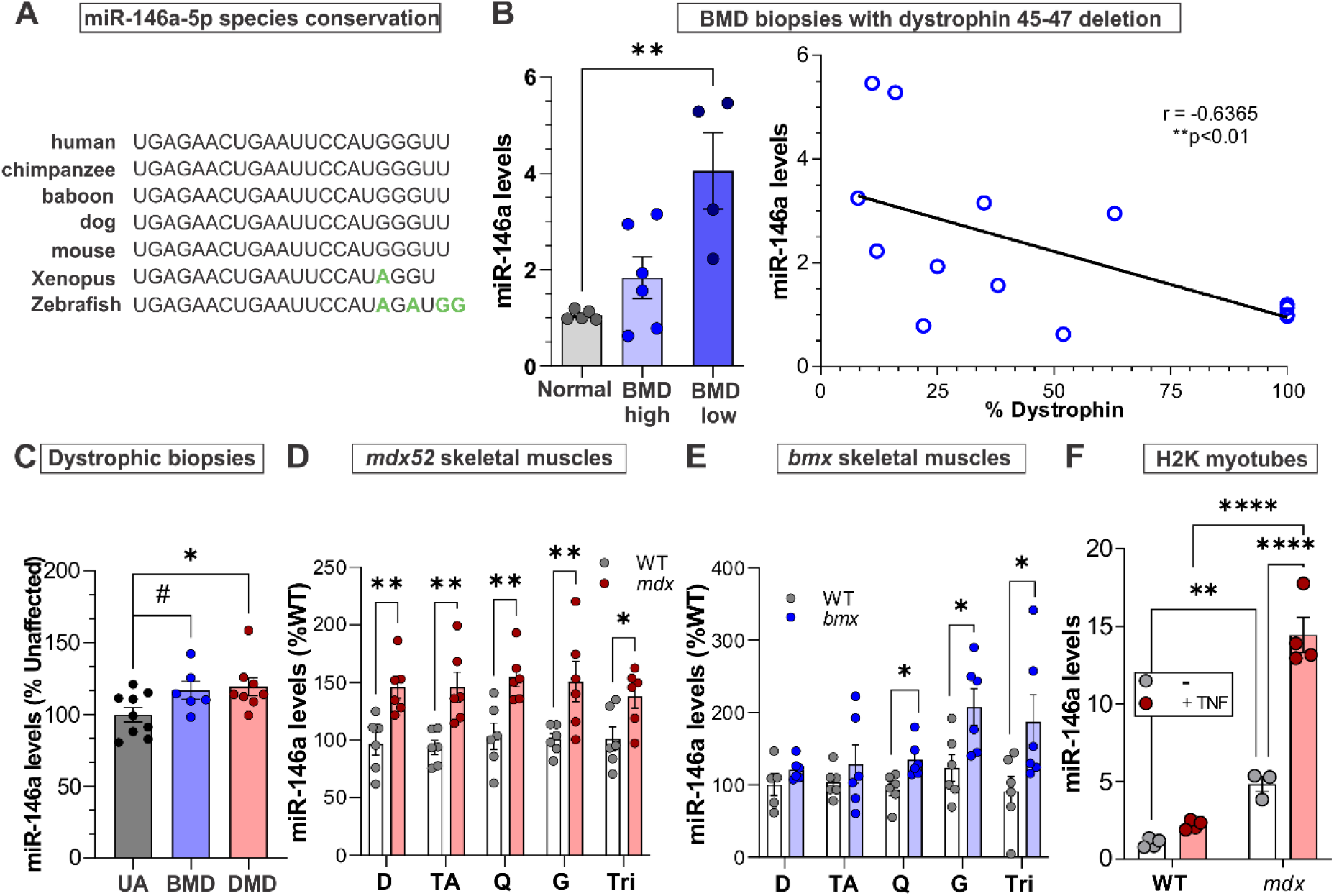
miR-146a is significantly elevated in BMD and DMD patients and model mice. (**A**) The sequence of miR-146a-5p is conserved across species. Note that in mammals there is 100% conservation. (**B**) Left; miR-146a levels are elevated in BMD patients with low dystrophin (<20% dystrophin of unaffected individuals). n=5 normal, 6 “BMD high” and 4 “BMD low”; ANOVA, **p<0.001, adapted from ^7^. Right; Dystrophin protein and miR-146a levels are inversely correlated in BMD patients. Spearman correlation, r=-0.6365, **p<0.01. Adapted from^7^. (**C**) miR-146a levels are significantly elevated in both BMD and DMD patients (taken from raw data in Eisenberg et. al^15^) n=9 unaffected (UA), 6 BMD and 8 DMD biopsies; ANOVA, #p<0.1, *p<0.05. (**D**) miR-146a levels are significantly increased in *mdx52* mouse diaphragm (D), tibialis anterior (TA), quadriceps (Q), gastrocnemius (G), and triceps (Tri). n=6/group; Two-way ANOVA, **p<0.01, *p<0.05. (**E**) miR-146a levels in *bmx* mouse diaphragm (D), tibialis anterior (TA), quadriceps (Q), gastrocnemius (G), and triceps (Tri). miR-146a is significantly increased in the latter 3 muscles. n=6/group, Two-way ANOVA, *p<0.05. (**F**) miR-146a is more highly expressed and induced in mdx vs. WT H2K myotubes treated with 10 ng/mL TNF-α (n=4 replicates per group) Student’s t-test, ****p<0.0001. Data are represented as mean ± S.E.M. See also Figure S1.

Previous studies have demonstrated that miR-146a is regulated by NF-κB in THP-1 macrophage cells^18^. We wanted to determine how induction of inflammation affects miR-146a levels in both *mdx* and wild-type (WT) myotubes (*mdx and WT H-2K*). Interestingly, in both WT and mdx myotubes TNF-α increases miR-146a, however this increase is only significant in *mdx* myotubes (**Figure 1F**). Analysis of qRT-PCR data of miR-146a also showed that 1) basal levels of miR-146a are significantly higher in *mdx* myotubes (373% increase over WT, p<0.01), 2) the overall induction of miR-146a in response to TNF-α is greater in mdx vs. WT (111% vs. 193% increase in miR-146a expression p<0.0001) and 3) the absolute levels of induced miR-146a were higher in *mdx* (555% increase, p<0.0001). We also observed high induction of miR-146a in *mdx* myotubes upon lipopolysaccharide (LPS)-induced inflammation (**Figure S1**). Overall, these findings show that miR-146a is upregulated in the muscle of BMD and DMD patients, as well as in mouse models of these diseases, and that dystrophic myotubes produce higher levels of miR-146a as compared to WT myotubes *in vitro*.

### miR-146a inhibits dystrophin translation

To validate our previous finding that miR-146a regulates dystrophin translation, we utilized the scanMiR toolkit^29^ to identify putative miR-146a binding sites or microRNA response elements (MREs) within the human dystrophin 3′UTR (**Figure 2A, B**). scanMiR identified multiple types of binding sites relative to the miR-146a seed sequence which is located at nucleotides 2-7 of the miRNA sequence^30^. These site types included a 7mer-m8 (perfect complementarity of the seed sequence to the UTR and complementarity extending to nucleotide 8), a 6mer-A1, (seed sequence complementarity at positions 3-7 and additional complementarity at nucleotide 8), a 6mer-A1 (seed sequences at positions 2-6 and an A at position 1), a wobbled 8mer (A at position 1, seed sequence complementarity interrupted by a single mismatch and additional complementarity at position 8), and several non-canonical binding sites^30^. In total, scanMiR predicted 17 binding sites, although only two showed strong dissociation constants (-log K_d_ <-3). The miR-146a MRE with the strongest predicted dissociation constant (log K_d_ -4.425) is located at position 1281-1288 of the 3′UTR (**Figure 2B, 2C**).

**Figure 2.**
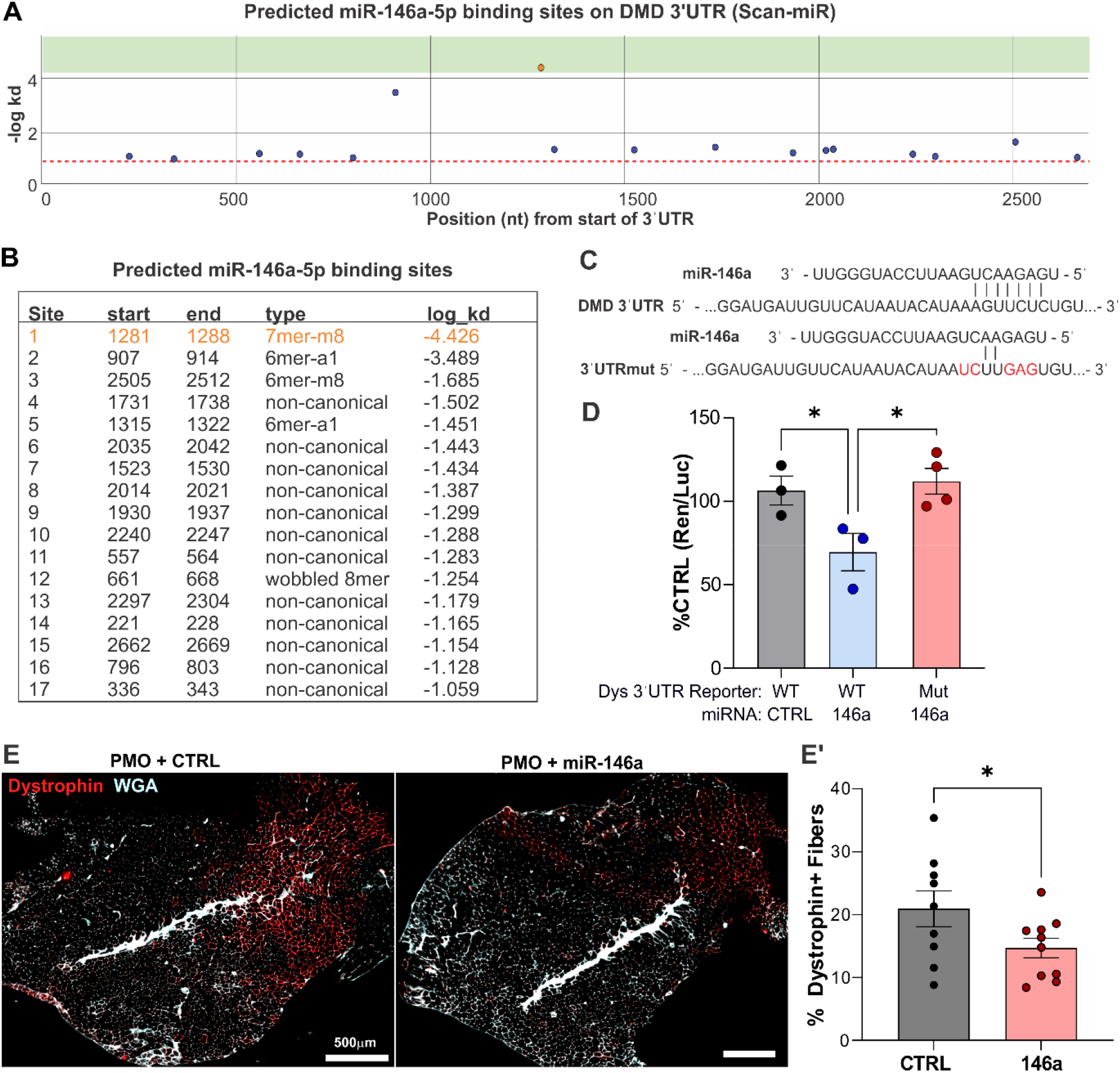
miR-146a targets the dystrophin 3′UTR and reduces exon skipping-mediated dystrophin restoration. (**A-B**) Predicted miR-146a-5p binding sites in the human *DMD* 3’UTR. (**A**) Scan-miR was used to search for putative miR-146a-5p binding sites within the human dystrophin 3′UTR sequence. The y axis denotes the dissociation constant (-log Kd), and the x axis denotes the location on the dystrophin 3′UTR in the 5′ to 3′ direction. (**B**) List of 17 putative miR-146a binding sites in the *DMD* 3’ UTR; orange denotes the strongest binding site and the one that was mutated in the luciferase construct in (**C**). The predicted binding sites include: a 7mer-m8 where there is perfect complementarity of the seed sequence with the UTR and this complementarity extends to nucleotide 8; a 6mer-A1 where there is seed sequence complementarity at nucleotide positions 3-7 and additional complementarity at nucleotide 8; a 6mer-A1, where the seed sequence shows complementarity to the UTR at positions 2-6 and additionally possesses an adenosine (A) at position 1; an wobbled 8mer, where there is an adenosine position 1, seed sequence complementarity at position 2-7 that is interrupted by a single mismatch and additional complementarity at position 8 and several non-canonical binding sites ^30^. (**C**) Schematic showing the binding site of miR-146a on the *DMD* 3’ UTR (top) and the base pairing of miR-146a with the *DMD* 3’ UTR when the binding site is mutated (bottom). (**D**) A luciferase reporter assay was used to examine miR-146a binding to the WT and mutant *DMD* 3’ UTR in HEK-293T cells. Transfection with miR-146a significantly reduces reporter activity in cells with WT *DMD* 3’ UTR whereas there is no reduction in reporter activity in cells with the mutated *DMD* 3’ UTR. n=3-4, ANOVA, *p<0.05. (**E**) The TAs of *mdx52* mice were injected with 2μg of exon skipping PMO and 10μg of either control or miR-146a. TA cross sections were stained for dystrophin (red) and wheat germ agglutinin (white). (**E’**) Quantification of dystrophin positive fibers; significantly fewer dystrophin-positive myofibers were observed in muscles co-injected with miR-146a compared to control. n=9-10, Student’s t-test, *p<0.05. Data represented as mean ± S.E.M. See also Figure S2.

Using this information, we next utilized a dystrophin 3′UTR luciferase reporter (WT) or a reporter where the 3’UTR MRE was mutated (Mut) to eliminate base-pairing of miR-146a at positions 2,3 and 6-8 (**Figure 2C**). WT and Mut constructs were transfected into HEK293T cells with either a control or miR-146a mimic sequence. Analogous to our previously reported results^7^, miR-146a significantly reduced WT 3′UTR luciferase expression while mutation of the miR-146a MRE attenuated this inhibition (**Figure 2D**). To support this *in vitro* data, we utilized the *mdx52* mouse, which is amenable to exon 51 skipping. We performed intramuscular tibialis anterior (TA) injections with 2 μg of an exon 51 skipping PMO and 10 μg of either a miR-146a mimic or a control sequence (n=10 per group). Fourteen days post-injection, muscles were harvested, and dystrophin immunofluorescence was performed. Using qRT-PCR, we observed significantly increased miR-146a in muscles co-injected with the miR-146a mimic sequence (**Figure S2**). Additionally, we found a significant decrease in dystrophin positive fibers in muscles co-injected with miR-146a as compared to muscle injected with a control sequence (36.1% decrease, p<0.05, **Figure 2E, E′**). Together these data suggest that miR-146a binds to the dystrophin 3′UTR to regulate its translation and high levels of miR-146a are inhibitory to successful dystrophin rescue via exon skipping.

### Generation of mdx/146a^-/-^ (146aX) mice

To investigate the potential therapeutic implications of miR-146a, we generated *mdx52* mice with body-wide deletion of miR-146a (*146aX*) by crossing several generations of *mdx52* and miR-*146a-/-* mice both *on* a C57/BL/6 background (**Figure 3A**). Deletion of miR-146a was determined by genotyping and validated by qRT-PCR in the gastrocnemius of 3-month-old mice (**Figure 3B**). The primary goal of generating these mice was to perform a proof-of-concept study to determine how deletion of miR-146a affects dystrophin restoration in *mdx52* mice via exon skipping. It has previously been shown that in dystrophin-deficient *mdx* mice with a nonsense mutation in exon 23 (*mdx23*), muscles contain sporadic clusters of dystrophin-expressing revertant fibers^31^. Revertant fibers occur due to spontaneous exon skipping (alternative splicing) and tend to accumulate as a function of age. Previous work has demonstrated that *mdx52* muscles have less revertant fibers as compared to *mdx23* mice^32^. We performed dystrophin immunofluorescence on muscle sections from wild-type, *mdx52* and *146aX* quadricep muscles to observe the baseline levels of revertant fibers in *mdx52* versus*146aX* muscles. As shown in **Figure 3C** there were minimal revertant fibers in both genotypes which totaled 0.16% in mdx and 0.14% of total fibers in *146aX* muscles (ns, *p*=0.6830). As typical high dose exon skipping experiments show 5-80% dystrophin positive fibers^33^ (depending on the specific muscle) the basal levels of revertant fibers observed in both *mdx* and 146aX here are negligible in comparison.

**Figure 3.**
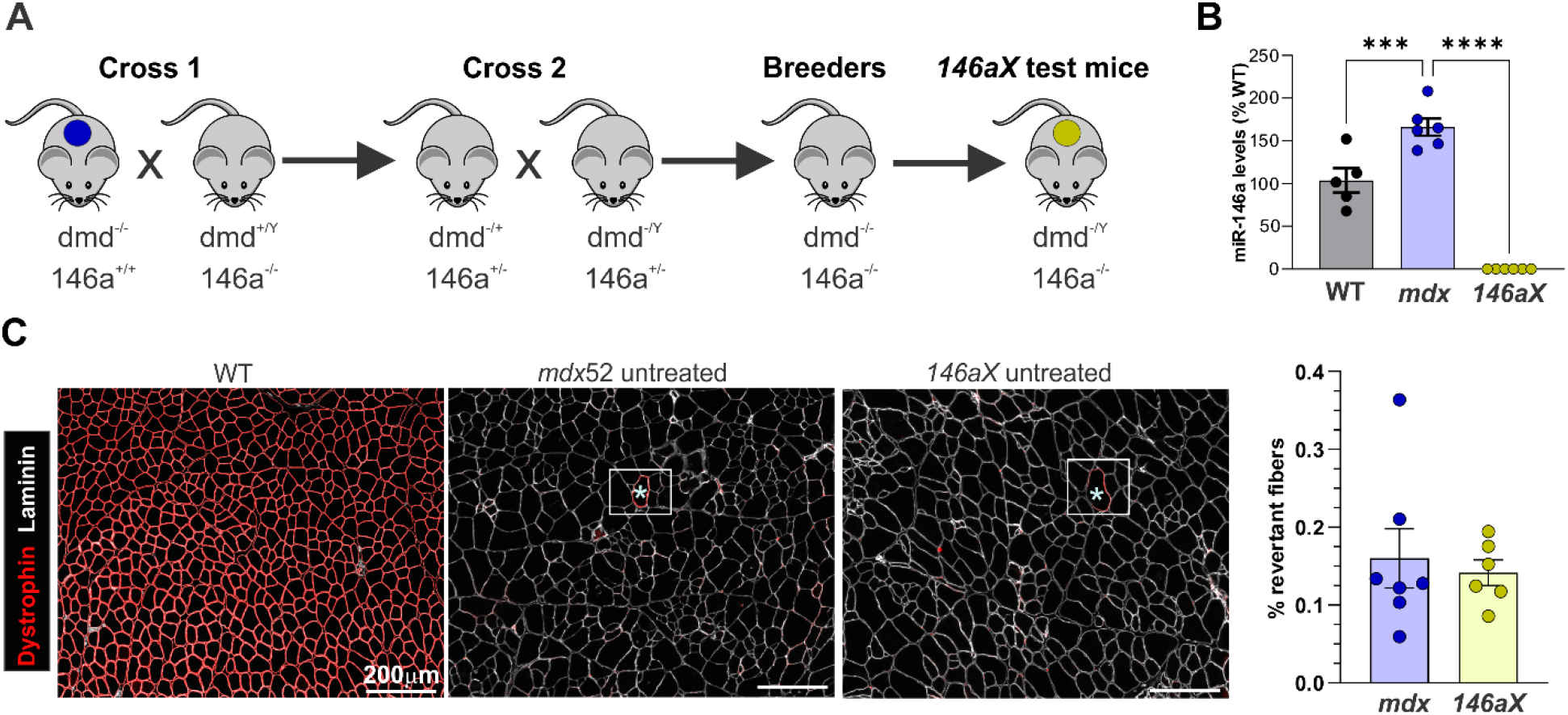
Generation of dmd^-/-^; 146a^-/-^ (*146aX* mice). (**A**) Breeding schematic to develop *146aX* mice. (**B**) qRT-PCR showing miR-146a levels in the gastrocnemius muscles of WT, *mdx*, and *146aX* mice. *146aX* mice have no detectable levels of miR-146a via qRT-PCR. n=6/group, ANOVA, ****p<0.0001. (**C**)Left; Representative dystrophin immunofluorescence (red) and laminin staining in *wild-type (WT)*, *mdx52* and *146aX* quadriceps showing strong dystrophin staining in WT muscle and no staining except for a few revertant fibers in *mdx52* and *146aX* muscles (white boxes with asterisks). Right; Quantification of revertant fibers in *mdx52* and *146aX* muscles shows that they represent only a small fraction of all fibers. There is no significant difference in revertant dystrophin-positive fibers between genotypes. n=6-7/group, students t-test, ns p=0.6830.

### Body-wide deletion of miR-146a increases dystrophin rescue after intramuscular PMO delivery

We next injected an exon 51 skipping PMO (2µg) into the TAs of *mdx52* and *146aX* mice (**Figure 4A**). Fourteen days post-injection, muscles were harvested and analyzed for extent of exon skipping and dystrophin protein. To analyze exon skipping we performed qRT-PCR on both *Dmd* exon 2 (total *Dmd* transcript levels) and the novel junction between *Dmd* exon 50 and 53 (skipped *Dmd* transcript levels) to determine the percent exon skipping achieved in each muscle as we previously described^34^. Levels of exon skipping ranged between 0.5% and 7%, and there were no significant differences observed in the extent of exon skipping when comparing *mdx52* and *146aX* TAs (p=0.48, **Figure 4B**). We next performed capillary Western immunoassays (Wes), as dystrophin quantification using this method has proven to be highly sensitive, reproducible, and quantitative over a large dynamic range^26^. While no differences in exon skipped *Dmd* transcript levels were observed, Wes protein quantification revealed a significant increase in dystrophin levels in *146aX* muscles (∼19% increase, p<0.05, **Figure 4C, C′**). Supporting this, dystrophin immunofluorescence showed a greater than 2-fold increase in dystrophin positive fibers in *146aX* TAs (p<0.01, **Figure 4D, D′**).

**Figure 4.**
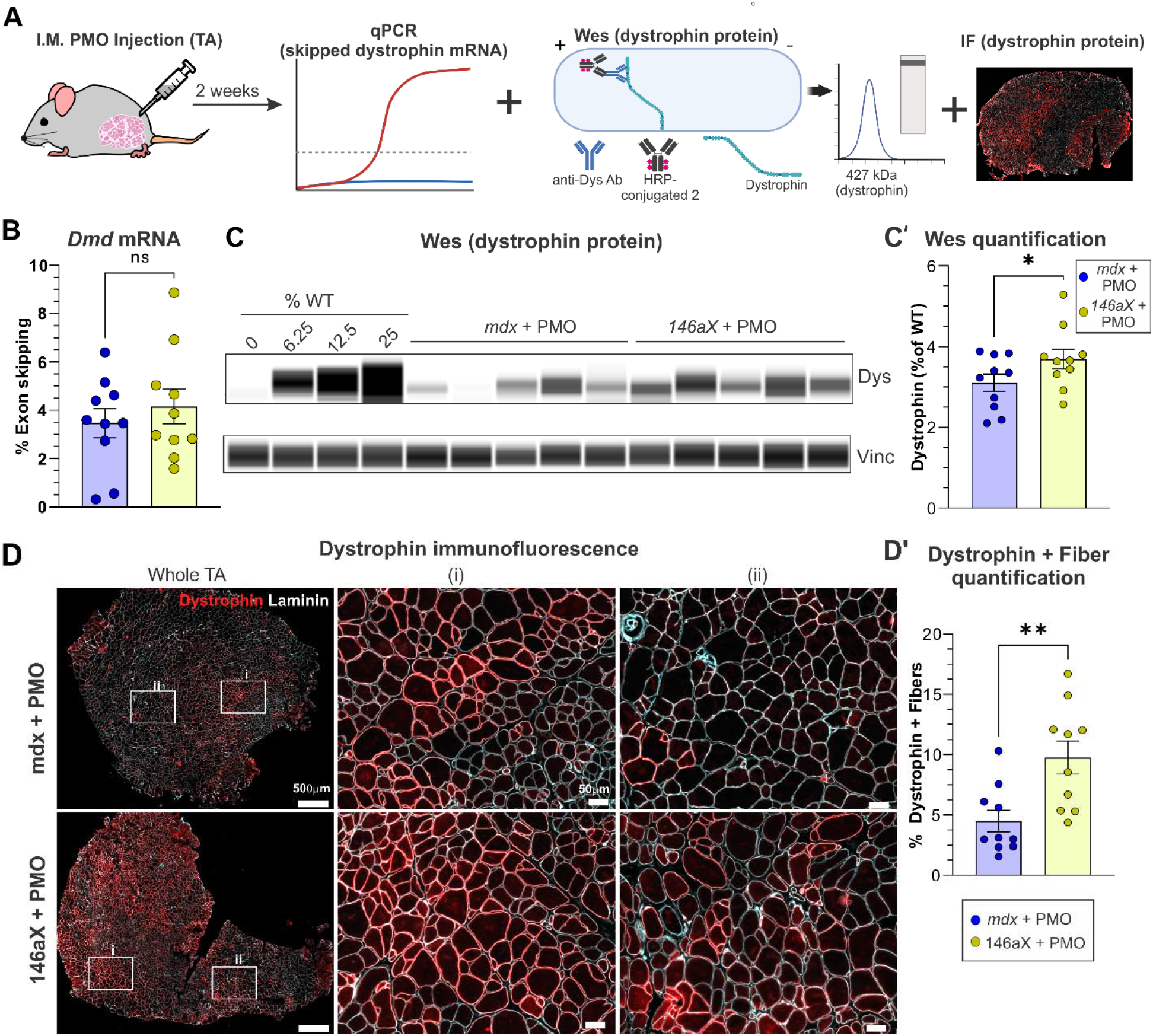
Deletion of miR-146a significantly increases dystrophin restoration in *mdx52* mice treated with intramuscular PMO. (**A**) Schematic of intramuscular PMO injection experimental design. *mdx* and *146aX* tibialis anterior muscles (TAs) were injected with 2µg of an exon 51 skipping PMO. 2 weeks post-injection muscles were harvested for analysis. n=10 muscles/group and 5 mice per group. Wes = Western capillary immunoassay, IF = immunofluorescence. (**B**) qRT-PCR was performed to quantify % exon skipping in PMO-injected *mdx52* and *146aX* TAs as previously described^34^. Exon skipping efficiency at the RNA level is not significantly between *mdx52* and *146aX* TAs. (**C**) Capillary-based immunoassay (Wes) was used to quantify levels of dystrophin protein. (**C’**) Dystrophin protein is significantly increased in *146aX* mice injected with PMO compared to *mdx* mice. (**D**) Immunofluorescence showing dystrophin staining in the TA of mdx and *146aX* mice with intramuscular PMO. (i) shows a zoomed in area of high dystrophin rescue and (ii) shows a zoomed in area of low dystrophin rescue. Bars = 500µm and 50 µm respectively. (**D’**) Quantification of dystrophin-positive myofibers. The percentage of dystrophin-positive myofibers following PMO injection is significantly increased in *146aX* mice compared to mdx mice. Images were blinded during quantification. Data represented as mean ± S.E.M. ns *p*>0.05, \**p*≤0.05, \*\**p*≤0.01, \*\*\*\**p*≤0.0001. See also, Figure S3.

### Body-wide deletion of miR-146a increases dystrophin rescue after systemic PMO treatment

We next performed systemic injections of a high dose PMO into 15-week-old *mdx52* or *146aX* mice (400 mg/kg) n=4 mice per group. One week later we performed a second high dose injection (400 mg/kg). Mice were phenotyped one week after the second dose of PMO and then sacrificed 2 weeks after the final dose. Diaphragm (Dia), tibialis anterior (TA), quadriceps (Quad), gastrocnemius (Gastroc), and Triceps (Tri) were collected for RNA and protein analysis (**Figure 5A**). When we analyzed the extent of exon skipping across muscles at the RNA level via qRT-PCR, no significant differences were observed between groups (**Figure 5B**), however, there was a non-significant trend towards increased exon skipping in the *mdx52* versus *146aX* muscles. Conversely, analysis of dystrophin protein levels via Wes showed significantly increased dystrophin in *146aX* muscles as compared to *mdx52* (∼55% increase, p<0.05, **Figure 5C, C′ Figure S3**). To account for differences in PMO delivery between muscles and genotypes, we also normalized dystrophin protein levels as measured by Wes, to exon skipping RNA levels measured by qRT-PCR. Interestingly, using this method to normalize dystrophin protein we found that on average, *146aX* muscles showed a 2.4-fold increase in dystrophin protein per “skipped” transcript (**Figure 5D**).

**Figure 5.**
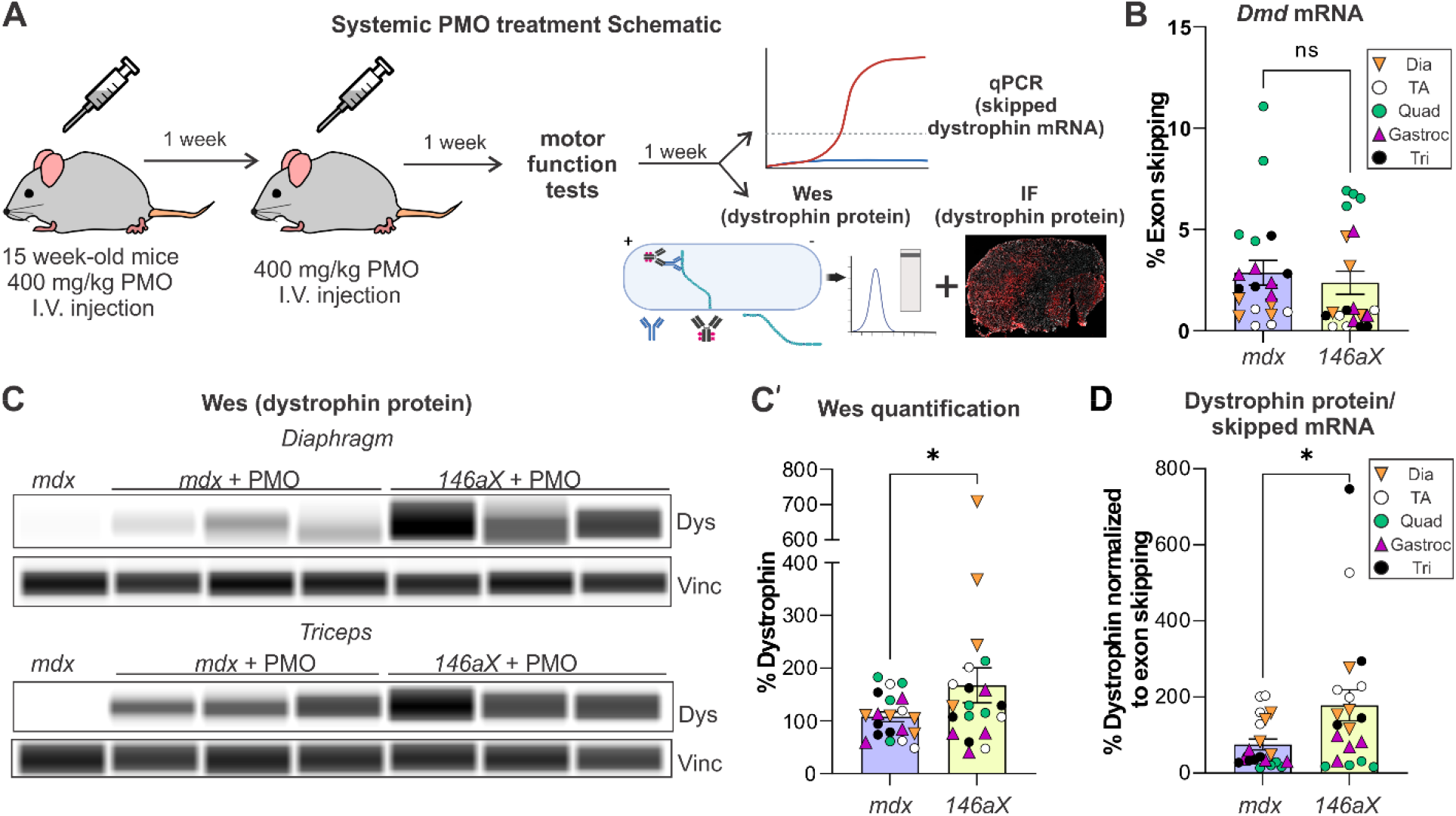
Deletion of miR-146a significantly increases dystrophin protein in *mdx52* mice treated with systemic PMO. (**A**) Schematic of experimental design for systemic PMO injections in *mdx* and *146aX* mice. *mdx52* and *146aX* mice were administered systemic PMO via the retroorbital sinus (400 mg/kg, exon 51 skipping PMO) at 15 weeks of age. One week later a second 400mg/kg PMO injection was performed. Muscle function tests were performed one week after the second injection and mice were sacrificed 2 weeks after the second injection. Muscles were analyzed by qRT-PCR to determine extent of skipped *Dmd* mRNA and by Western capillary immunoassay (Wes) and immunofluorescence (IF) to determine dystrophin protein levels (IF). n=4 mice and 5 muscles per group. (**B**) Exon skipping efficiency at the RNA level via systemic injection of PMO in *mdx* and *146aX* mice is not significantly different. Student’s t-test. (**C**) Capillary-based immunoassay (Wes) was used to quantify dystrophin protein levels in skeletal muscle of *mdx* and *146aX* mice systemically treated with exon-skipping PMO via retroorbital injection. Depicted is a virtual Wes blot. (**C’**) Wes quantification of dystrophin protein levels in skeletal muscle. Dystrophin protein is significantly increased in *146aX* mice versus *mdx52* mice. Student’s t-test, *p<0.05. Note: % Dystrophin was calculated by normalizing to the average *mdx* intensity for each muscle and setting it to 100%. (**D**) Dystrophin protein levels from Wes normalized to % *Dmd* exon skipped transcripts. There is a higher ratio of dystrophin restoration in *146aX* mice compared to *mdx* mice. Data represented as mean ± S.E.M. Student’s t-test, *p<0.05.

To support this data, we additionally performed quantification of dystrophin positive fibers using immunofluorescence analysis. Analysis showed a greater dystrophin restoration via an increase in dystrophin positive fibers in *146aX* muscles (43.5% increase p<0.05, **Figure 6A, A′, Figure S4**). This difference persisted after normalizing to the extent of skipped *Dmd* transcripts, showing a 3.5-fold increase in dystrophin positive fibers per “skipped” transcript (**Figure 6B**). In our analysis we observed high variability in the “brightness” or intensity of dystrophin positive fibers. To investigate this further we generated a visual standard to label the intensity of dystrophin positive fibers as either “high” or “low” (**Figure 6C**) and then quantified each group. Interestingly, we found that while there was a modest increase in the extent of “high” intensity fibers in *146aX* muscle (35.6% increase, p<0.1), “low” intensity fibers were 60.52% higher in *146aX* muscles (p<0.05, **Figure 6D**). A potential interpretation of this difference is that the absence of miR-146 stabilizes “skipped” dystrophin transcripts and more protein continues to be produced from these skipped transcripts. Collectively, this data supports the idea that reducing miR-146a can markedly improve exon-skipping-mediated dystrophin restoration in dystrophin-deficient mice.

**Figure 6.**
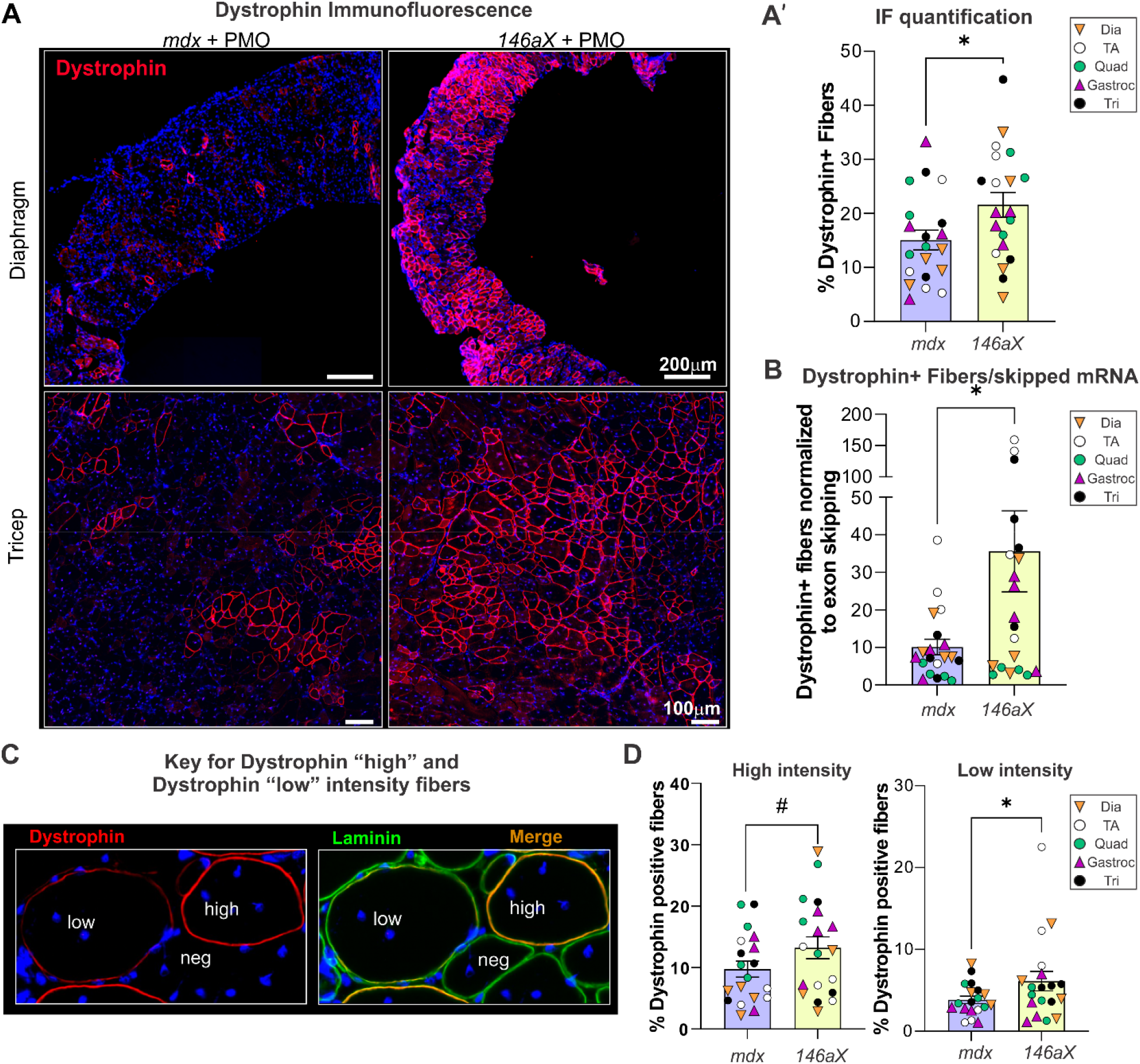
Deletion of miR-146a significantly increases dystrophin positive fibers in *mdx52* mice. (**A**) Dystrophin immunofluorescence (red) in PMO-injected *mdx* and *146aX* diaphragm and triceps muscles. Bar= 200mM and 100 mM, respectively. Muscles were counterstained with DAPI (nuclei) and laminin (green, not shown) to count total muscle fibers). (**A’**) Quantification of dystrophin-positive myofibers, (n=4 mice and 5 muscles per group). The percentage of dystrophin-positive myofibers following PMO injection is significantly increased in *146aX* mice compared to mdx mice. (**B**)The percentage of dystrophin positive fibers normalized to the percentage of *Dmd* exon skipped transcripts. There is a higher ratio of dystrophin restoration in *146aX* mice compared to *mdx* mice. Data represented as mean ± S.E.M Student’s t-test, *p<0.05. (**C**) Visual standard for classification of dystrophin high, low, and negative fibers. (**D**) Left; Quantification of “high” intensity dystrophin-positive fibers. Right; quantification of “low” intensity dystrophin positive fibers. Images were blinded during quantification. Student’s t-test, #p<0.1, *p<0.05. All data represented as mean ± S.E.M. Dia, diaphragm; TA, tibialis anterior; Quad, quadriceps; Gastroc, gastrocnemius; Tri, triceps. See also Figure S4.

### Improved muscle function in PMO-treated 146aX

We performed functional testing on 17-week-old, treated *mdx52* and *146aX* mice (1 week after second PMO injection). *146aX* mice showed a 24.4% increase in hindlimb grip strength versus *mdx52* mice (p<0.05, **Figure 7A**), while forelimb grip strength showed similar measurements for treated *mdx52* and *146aX* (**Figure S5A**). Suspension time in the two-limb wire hang in treated *146aX* mice showed an increase (39.4% increase) but did not reach statistical significance (p=0.163, **Figure 7A**).

**Figure 7.**
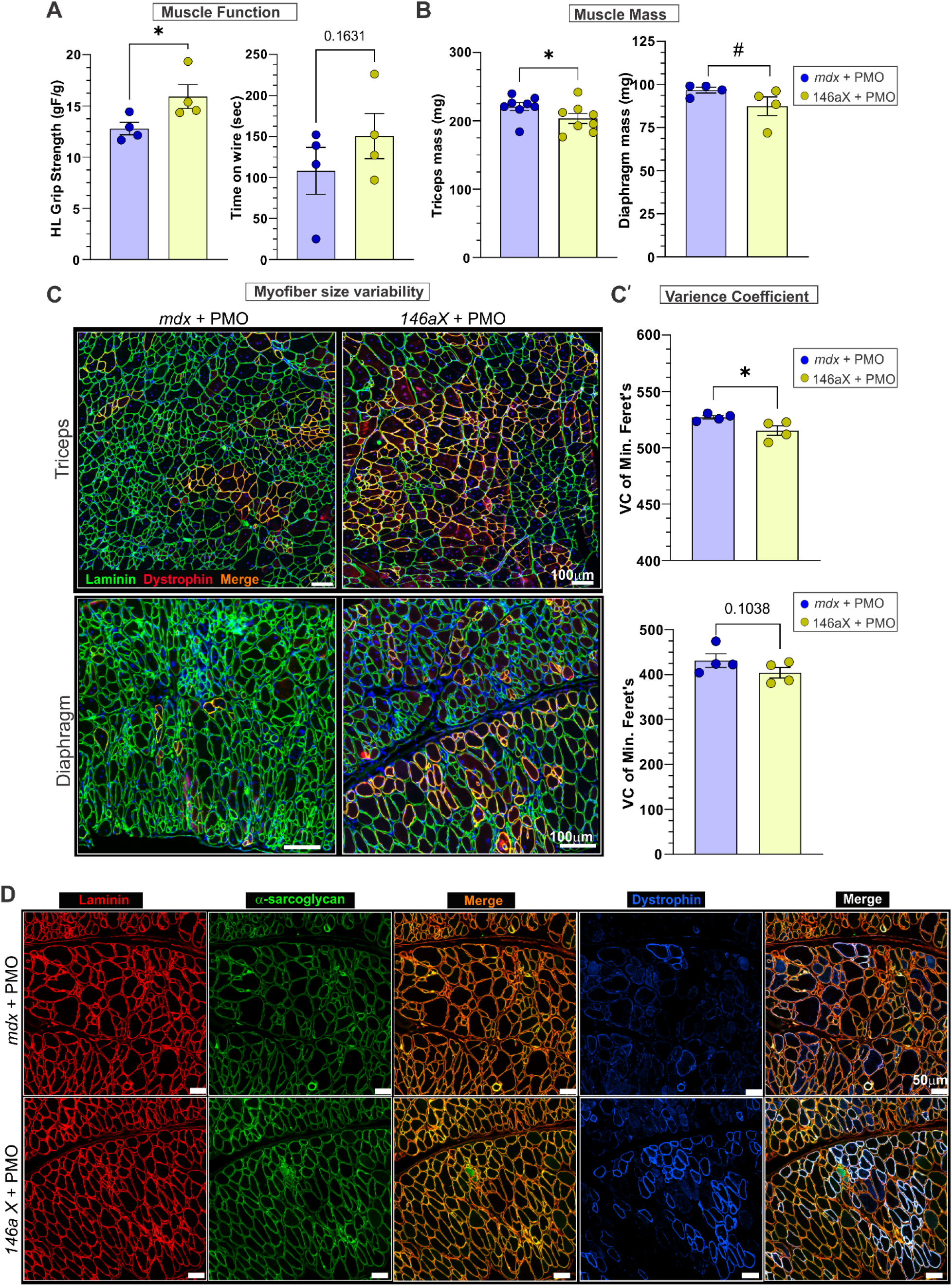
Functional and histological analysis of PMO-treated *mdx52* and *146aX* mice. (**A**) Muscle function tests in *mdx52* and *146aX* mice. Left; Hindlimb grip strength was performed and normalized to body weight. Right; Wire hang-time assay. n=4 mice per group; Student’s t-test. (**B**) Terminal tissue masses of triceps (n=8) and diaphragms (n=4). Student’s t-test, #p<0.1, *p<0.05. All data represented as mean ± S.E.M.(**C**) Immunofluorescence of muscle sections using laminin (green), dystrophin (red) and DAPI (blue) staining of treated *mdx52* versus *146aX* muscles to visualize both dystrophin restoration and myofiber size. Bar=100µm. (**C’**) Quantification of myofiber size variability via the variance coefficient of minimal Feret’s diameter. (**D**) To visualize how dystrophin restoration affects localization of a dystrophin-associated protein (α-sarcoglycan), laminin (red), a-sarcoglycan (green), and dystrophin (blue) were visualized via immunofluorescence in *mdx52* and *146aX* diaphragms. Note that α-sarcoglycan is brighter in treated *146aX* diaphragms which is apparent by the yellow laminin/a-sarcoglycan co-localization (merge) image versus the more orange hue in the laminin/α-sarcoglycan image in treated *mdx52* diaphragms. Bar = 50µm. See also Figure S5.

### Reduced pathology in treated 146aX muscles

Given the increases we saw in dystrophin rescue in *146aX*, most notably in the diaphragm and triceps muscles, we investigated the therapeutic benefit of systemic PMO treatment on these muscles by performing additional endpoint analysis on tissues. In *mdx* mice, limb muscles undergo dramatic necrosis, which is followed by regeneration, increased hypertrophy and increased muscle mass^35^. Analyzing tissue weights, the mass of both triceps and diaphragm muscles showed reductions in *146aX* mice (**Figure 7B**), indicating reduced pseudohypertrophy. The mass of gastrocnemius muscle was also significantly reduced, while other muscles showed non-significant reductions in *146aX* versus *mdx52* genotypes (**Figure S5B**). We observed no marked differences in endpoint body mass between treated *mdx52* and 146aX mice, however, triceps showed significant reduction in *146aX* mice when normalizing to body weight, while other muscles showed non-significant reductions (**Figure S5C,D**). Additionally, laminin staining showed visually less myofiber size variability in diaphragm and triceps muscles (**Figure 7C**) and analysis of minimal Feret’s diameter in laminin-stained muscle confirmed this reduction (variance coefficient, **Figure 7C′**) consistent with reduced asynchronous regeneration and a “healthier” muscle state in treated *146aX*^36^. These data were corroborated with hematoxylin and eosin (H&E) analysis which showed visually healthier muscle in treated *146aX* diaphragms muscles; quantification showed reduced inflammation and necrosis in these muscles (**Figure S5E**). Additionally, immunofluorescence analyses of the mouse macrophage marker F4/80 showed visually reduced macrophage infiltration in treated *146aX* diaphragms (**Figure S5F**). We additionally analyzed the dystrophin associated protein, α-sarcoglycan in treated diaphragms. Interestingly, we observed stronger α-sarcoglycan staining in treated *146aX* diaphragms visible as increased yellow areas of α-sarcoglycan (green) and laminin (red) co-localization, and this staining was especially pronounced in myofibers with dystrophin positivity (**Figure 7D**)

## Discussion

In the study presented here, we find miR-146a inhibits dystrophin protein levels *in vitro* and *in vivo* and is induced by inflammation in muscle. We also find that genetic deletion of miR-146a increases local dystrophin protein levels after intramuscular injection of an exon 51 skipping PMO in *mdx52* mice and increases body-wide dystrophin protein levels after systemic injection. Increased dystrophin rescue in systemically treated *146aX* mice also corresponds to increased muscle function, reduced histopathology and reduced myofiber size variability. To our knowledge, this is the first report showing that genetic deletion of a single miRNA can increase dystrophin protein in combination with administration of an exon skipping PMO.

We show here that PMO-treated *146aX* mice exhibit increased dystrophin protein levels compared to *mdx52*, while the extent of skipped *Dmd* transcript levels remains similar between groups. The observed increase in dystrophin protein but not dystrophin transcript levels is in line with our previous data in BMD biopsies and in BMD model mice showing that, while dystrophin transcript levels are the same, dystrophin protein levels are significantly reduced and negatively correlate with levels of miR-146a and other DTMs^7, 8^, and that most often miRNAs inhibit translation but do not promote mRNA decay^17^.

In our previous work, we identified and characterized 7 miRNAs that are elevated in BMD muscle biopsies, are inversely correlated with dystrophin protein levels, and found that these miRNAs target the dystrophin 3′UTR *in vitro*^7^. We collectively refer to this group of miRNAs as dystrophin targeting miRNAs or DTMs. This previous report also showed that treatment with the corticosteroid prednisone or the dissociative steroid vamorolone reduced expression of 3 DTMs including miR-146a in *mdx* mice^7^. In a separate report, we demonstrated that administration of either prednisone or vamorolone to low-dystrophin expressing BMD model (*bmx*) mice reduces expression of miR-146a and other DTMs (miR-31, miR-146b, miR-223) and increases dystrophin levels by 50%^21^. The goal of the previous report was to determine if reducing expression of multiple miRNAs could increase dystrophin protein levels. In the present study, our goal was to determine if targeting a single dystrophin-targeting miRNA could improve dystrophin restoration via exon skipping PMOs.

By using a genetic deletion model to ablate miR-146a in *mdx52* mice, this initial study provides proof-of-principal that targeting miR-146a could potentially lead to increased dystrophin rescue in DMD exon skipping. Upon systemic delivery of PMO, we found increased dystrophin protein levels in dystrophic mice lacking miR-146a. Specifically, Wes immunoassay quantification showed a 56.6% increase in dystrophin levels in *146aX* muscles. Additionally, using dystrophin immunofluorescence, we show that extent of dystrophin positive fibers increases from ∼15 to 25% in 146aX muscles and that on average, deletion of miR-146a enable ∼2.5 to 3.5-fold more dystrophin protein to be expressed from each “skipped transcript.” Together these data suggest that genetic deletion of miR-146a results in more efficient dystrophin protein production. We observed more myofibers expressing a “low” level of dystrophin via immunofluorescence. This could potentially be explained by the fact that mice were sacrificed two weeks after the final PMO injection and that fibers with “low” dystrophin were just starting to accumulate sufficient dystrophin protein from skipped transcripts. Potentially, if mice have been sacrificed 4-6 weeks after the final PMO injection, it would have allowed more time for dystrophin protein to accumulate from “skipped” transcripts which would have yielded a greater difference between *mdx52* and *146aX* genotypes. Future studies will determine how miR-146a deficiency affects the extent of *mdx52* muscle turnover, the persistence of skipped transcripts and the overall accumulation of dystrophin protein after PMO administration.

Our data show that deletion of miR-146a results in significant increases in dystrophin protein as measured via Wes and immunofluorescence. The extent of dystrophin rescue reported from DMD exon skipping clinical trials for Eteplirsen, golodirsen and casimersen is, on average, about 0.9-1.9% of unaffected levels^10, 13^ (NCT02500381). The target amount of dystrophin restoration needed to show a functional benefit in DMD is still unclear. Some reports using severe mouse models show 4% dystrophin is sufficient for improved muscle function and survival ^37–39^. Other reports show that 20% dystrophin improves symptoms^40^ and 15% could be sufficient to protect against contraction-induced injury in DMD model mice but more than 40% dystrophin is needed to also improve muscle force^41^. Additionally, we previously reported that even in BMD model (*bmx*) mice that express up to 30-50% dystrophin, muscles show susceptibility to contraction-induced injury and have reduced function as measured by grip strength and wire hang measures^8^. The most recent clinical data for the NS Pharma drug Viltolarsen shows more promising preliminary data, with 5.9% dystrophin measured in muscle biopsies^42^. Even with the increased efficacy of viltolarsen, improvement in the extent of dystrophin restoration will be important to yield the full potential of exon skipping therapeutics in DMD. Our study here suggests that targeting miR-146a could potentially contribute to that overall goal. We should caveat this statement, however, by noting that our proof-of-concept study here is preliminary and that additional pre-clinical studies to target miR-146a and to measure potential side effects of miR-146a inhibition are required before determining the true clinical translatability.

Cacchiarelli et. al^16^ was the first to show that a miRNA that is significantly upregulated in *mdx* muscles, miR-31, could target the dystrophin 3′UTR and inhibit dystrophin protein production. Our lab later built on this data using BMD del45-47 biopsies and described seven additional miRNAs that target dystrophin, including miR-146a. Following the initial miR-31 publication, Wells and Hildyard performed intraperitoneal injections of an exon 23 skipping vivo-PMO alone or in combination with a miR-31 protector sequence (to prevent miR-31 binding to the dystrophin 3’UTR) or a “scrambled protector” sequence^43^. While they observed increased dystrophin in cell culture experiments by inhibiting or blocking miR-31, this result did not translate *in vivo*^43^. There were, however, limitations to this study as the authors discuss. Specifically, they postulated that delivery of the vivo-morpholinos overall was low, with almost no delivery to hindlimbs, modest delivery to the diaphragm and less to the abdominal wall, and therefore it wasn’t clear whether the miR-31 target blocking strategy failed or the co-delivery of the exon skipping PMO, and target blocker was too low to observe any measurable differences in dystrophin protein. Based on their report and interpretation, here, we chose to use a genetic approach to ablate expression of miR-146a so that the efficiency of delivery of an inhibitor or blocking sequence was not a confounding factor in analysis. Moving forward we plan on performing studies where we will conjugate a miR-146a inhibitor to a cell penetrating peptide or antibody to determine how co-delivery of a miR-146a inhibitor or target blocking sequence with an exon skipping PMO affects dystrophin protein restoration in dystrophic mice.

Elevated levels of miR-146a have been described in numerous other chronic inflammatory and muscle disorders including myositis^44^, Miyoshi myopathy^15^, limb girdle muscular dystrophy^15^, FSHD^45^, rheumatoid arthritis^46^, Sjogren’s syndrome^47^, inflammatory bowel disease^19, 48, 49^, and asthma^50^ suggesting it plays a role in diseases where inflammation persists. Previous studies have reported that miR-146a is rapidly elevated in response to inflammatory stimuli, but then “turns off” the NF-κB signaling pathway via binding to the *Traf6* and *Irak1* 3′UTRs^18, 51, 52^. Because of this, it is generally regarded as “anti-inflammatory.” However, more recent work has described a mechanism where exogenous miR-146a-5p, but not its duplex precursor, is “pro-inflammatory” and induces inflammation via activation of TLR7 while it’s duplex precursor functions as part of a negative feedback loop to “turn off” NF-κB^53^. The work of Wang et. al described above therefore suggest that the narrow definition of a miRNA such as miR-146a as either anti-or pro-inflammatory is perhaps too reductive and more work is required to fully delineate the comprehensive role of each microRNA in both a healthy and a disease state.

It should be noted that another group also generated *mdx:miR-146a-/-* mice^54^, however, the overall aim of that work was significantly different from the work we present here. The previous study sought to test the hypothesis that miR-146a is anti-inflammatory and that genetic deletion of miR-146a in *mdx* would worsen the natural etiology of disease; after analysis they reported no signficant differences between the *mdx* mice with or without miR-146a expression. The data we present here demonstrates that miR-146a deletion benefits dystrophin rescue in *mdx52* mice treated with an exon 51 skipping PMO.

Interestingly, we found elevated miR-146a in murine myositis muscle and human muscle biopsies of inclusion body myositis^44^. In these same muscles, we observed reduced dystrophin staining via Western blot as well as reduced and discontinuous dystrophin staining^44^. Additionally, another group recently reported reduced dystrophin and increased dystrophin-targeting microRNAs (miR-146a, miR-223) in immune-mediated necrotizing myopathy^55^. Thus, it is possible that DTMs such as miR-146a contribute to a secondary dystrophin deficiency that would impair overall muscle function and the therapeutic targeting of this inflammation-regulated miRNA could be beneficial in the context of other muscle disorders.

One potential way to improve DMD exon skipping outcomes is to harness miRNA therapeutics as a combination therapy with the goal of inhibiting miR-146a-5p levels to enable increased dystrophin protein production from skipped dystrophin transcripts. This strategy could also potentially be applied as a stand-alone therapy in BMD where lower-than-normal levels of dystrophin are observed in muscle. There are a few different ways in which miR-146a inhibition or reduction can be achieved. These include the use of 1) a miR-146a inhibitor (anti-miR), 2) an oligonucleotide that prevents miR-146a from binding to the dystrophin 3′UTR (target blockers or blockmiRs), or 3) a small molecule inhibitor. Currently, there are several anti-miR therapeutics in clinical trials that show promising preliminary results (reviewed in^56^) including CDR132L, a locked nucleic acid (LNA) targeting miR-132 in heart failure^57^, Miravirsen/SPC3649, an LNA targeting miR-122 in patients with chronic hepatitis C virus^58^, and MRG-110, an LNA targeting miR-92a to benefit wound repair failure^59^. Additionally myotonic dystrophy (DM1) preclinical studies, using either a miR-23b antagomiR^60^ or a peptide-conjugated blockmiR to prevent miR-23 binding to MBNL^61^, have shown these strategies improve muscle function in mouse models. We have also observed reduced levels of miR-146a through treatment with anti-inflammatory small molecules, either vamorolone or prednisone, in *mdx* mice^7, 20^. A recent report from our lab shows that while both prednisone and vamorolone reduce miR-146a and increase dystrophin in the gastrocnemius muscle of *bmx* mice, only vamorolone significantly increases dystrophin levels in the heart^21^. While corticosteroids will likely always be given as a combination therapy along with exon skipping PMOs or other dystrophin restoration therapies, pre-clinical studies that investigate which corticosteroids (prednisone vs. deflazacort vs. vamorolone) yield the greatest increases in dystrophin restoration will be informative to current and future DMD treatment regimens.

In conclusion, the data presented here shows that genetic deletion of miR-146a in *mdx52* mice increases levels of therapeutic dystrophin protein restoration without affecting the extent of skipped dystrophin transcript levels. These data provide a strong rationale for investigating how therapeutic targeting of miR-146a and other DTMs may impact dystrophin protein levels in DMD exon skipping, as well as in other muscle diseases more broadly.

## Author Contributions

NM contributed to experiments, data analysis, manuscript writing and interpretation; KC performed data analysis; CS performed F4/80 immunofluorescence, CT performed experiments and analysis, CH contributed to experimental design, writing and interpretation, AF contributed to experimental design, experiments, analysis, and manuscript writing.

## Acknowledgements

We would like to thank Dr. Shin’ichi Takeda for the generous gift of the *mdx52* mice.

