## Supplemental Figures and Legends for "Deletion of miR-146a enhances therapeutic protein restoration in model of dystrophin exon skipping"

***Supplementary Material***

**Deletion of miR-146a enhances**

**therapeutic protein restoration in**

**model of dystrophin exon skipping**

**Nikki M. McCormack^1^, Kelsey A. Calabrese^1^, Christina M. Sun^1^, Christopher B. Tully^1^, Christopher R. Heier^1,2^, Alyson A. Fiorillo^1,2^**

^1^Center for Genetic Medicine Research, Children’s National Hospital, Washington, District of Columbia, USA

^2^Department of Genomics and Precision Medicine, George Washington University School of Medicine and Health Sciences, Washington, District of Columbia, USA

***Correspondence:**

Alyson A. Fiorillo, PhD

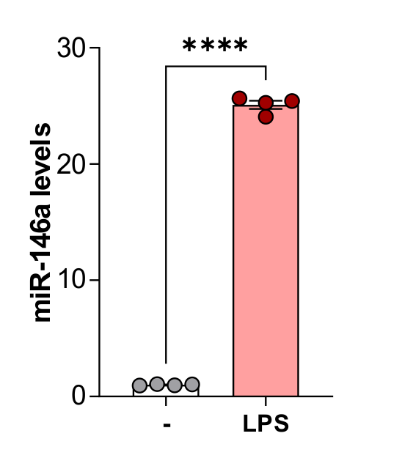

**Figure S1.miR-146a is induced by inflammation.** qRT-PCR shows miR-146a is significantly elevated in *mdx* H2K myotubes treated with LPS for 24 h. (n=4 replicates) Student’s t-test, ****p<0.0001. Data represented as mean ± S.E.M.

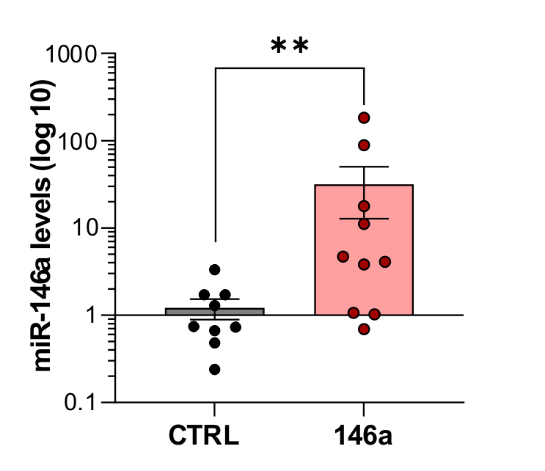

**Figure S2 miR-146a is increased in injected muscles.** qRT-PCR confirms miR-146a levels are significantly increased in TAs injected with miR-146a. n=9-10, Student’s t-test, **p<0.01. Data represented as mean ± S.E.M.

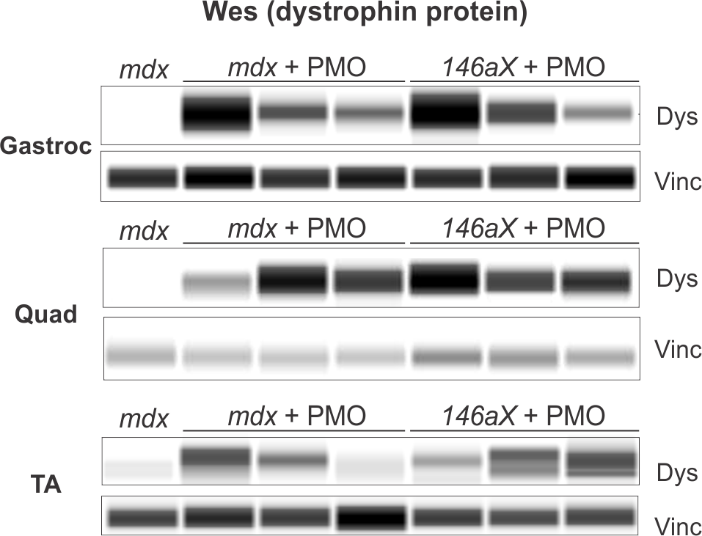

**Figure S3. Deletion of miR-146a significantly increases dystrophin restoration in *mdx52* mice treated with systemic PMO.** Capillary-based immunoassay (Wes) was used to quantify dystrophin protein levels in skeletal muscle of *mdx* and *146aX* mice systemically treated with exon-skipping PMO via retroorbital injection. Depicted is a virtual Wes blot for gastroc, quad and TA muscles. Dys, dystrophin; Vinc, vinculin

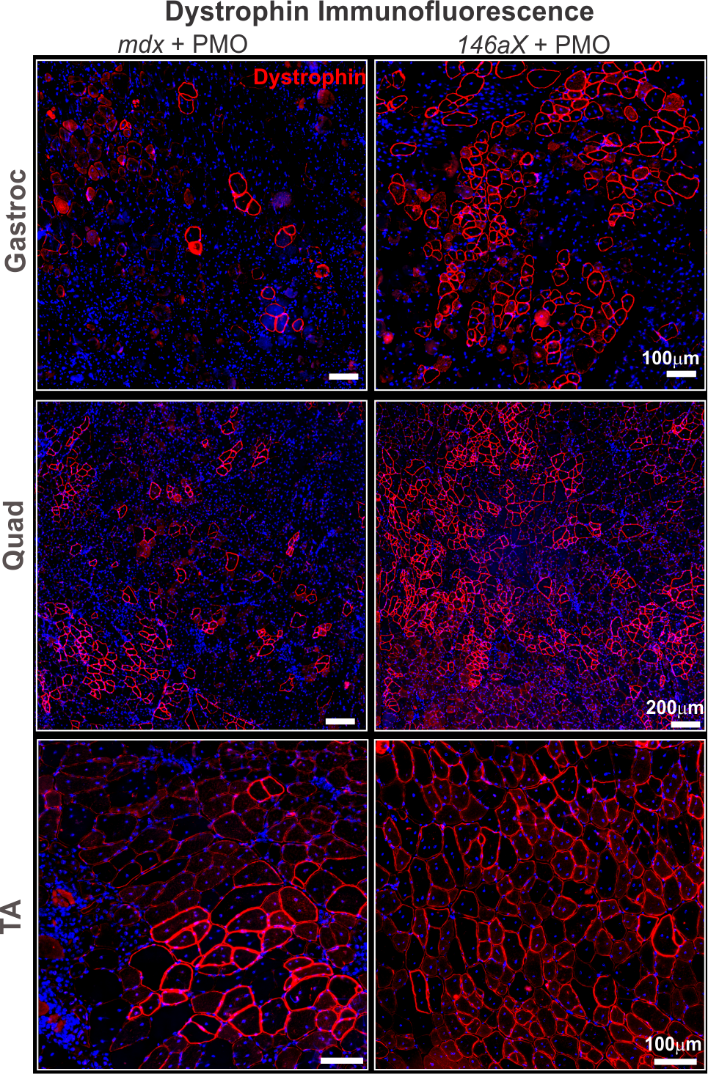

**Figure S4. Deletion of miR-146a significantly increases dystrophin positive fibers in *mdx52* mice.** Dystrophin immunofluorescence (red) in PMO-injected *mdx52* and *146aX* gastroc, quad and TA muscles Bar= 100µm (gastroc) and 200µm (quad) and 100µm (TA). Muscles were counter-stained with DAPI.

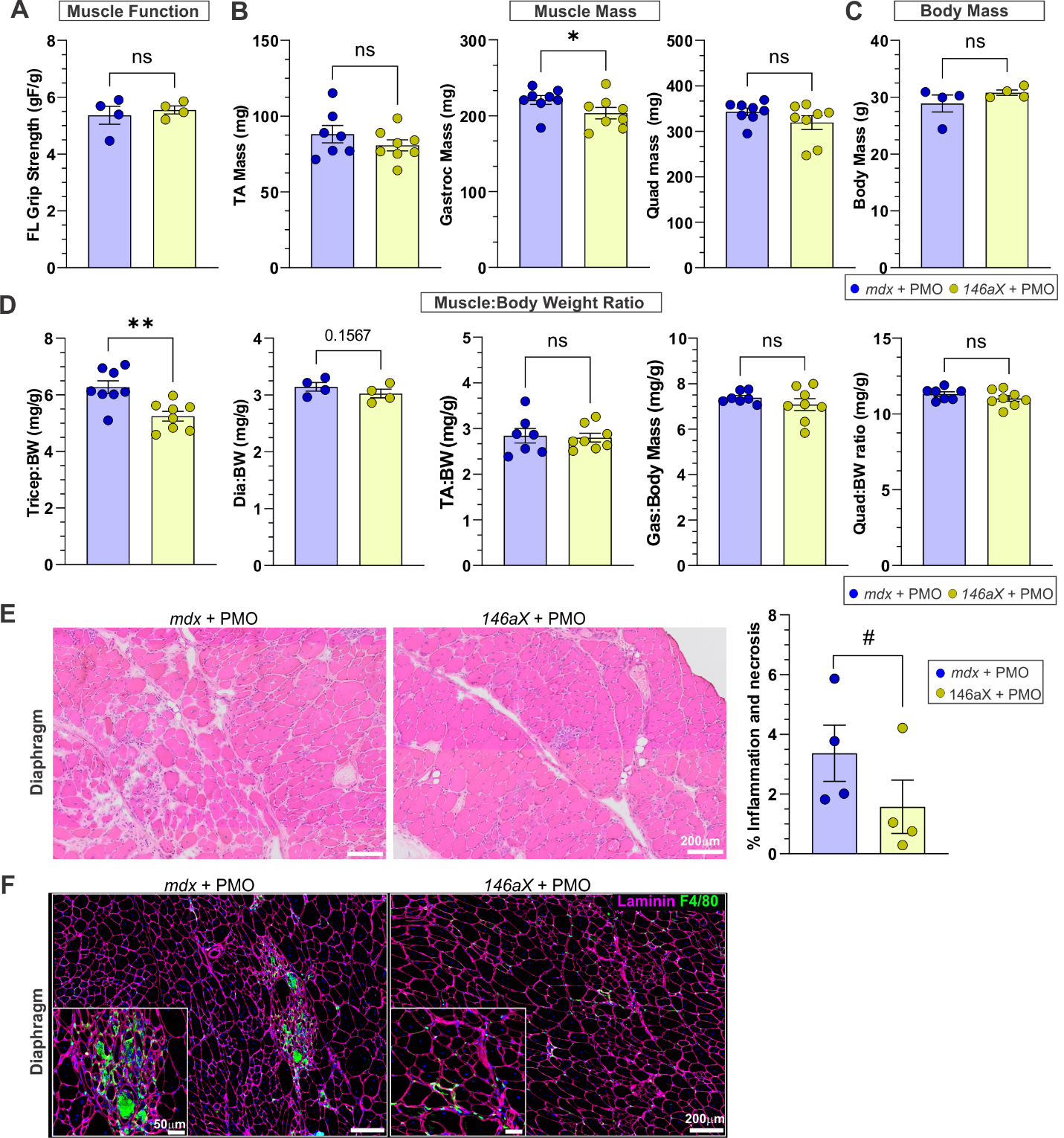

**Figure S5. Functional and histological analysis of PMO-treated *mdx52* and *146aX* mice.** (A) Forelimb grip strength measurements. n=4. (B) Terminal tissue weights of tibialis anterior (TA), gastrocnemius (gastroc) and quadriceps (quad) muscles. n=8. (C) Terminal body mass of treated mice. n=4. (D) Indicated tissue weights normalized to body mass (n=8 Tri, TA, Gas, Quad; n=4 dia). (**E**) Hematoxylin and eosin-stained diaphragm muscles from treated *mdx52* or *146aX* mice. *Left*; representative images, right: quantification of % inflammation and necrosis in treated muscles *n* = 4. (F) Macrophage (F4/80, green) and laminin (pink) staining in treated *mdx52* and *146aX* diaphragm muscles showing reduced presence of infiltrating macrophages in *146aX*. Student’s t-test, #p<0.1, *p<0.05. All data represented as mean ± S.E.M.
